# Comparative wastewater virome analysis with different enrichment methods

**DOI:** 10.1101/2025.03.25.645222

**Authors:** Matthew Thornton, Gabriela Eder, Fabian Amman, Anastasiia Pantielieieva, Julia Vierheilig, Andreas Bergthaler

## Abstract

Wastewater-based epidemiology (WBE) has proven its value for public health. Physical concentration of virus particles is a crucial step for WBE to permit a sensitive and unbiased characterization of the catchment virome. Here we evaluate five different virion concentration techniques, including polyethylene glycol precipitation (PEG), vacuum-based direct capture (VDC), ultrafiltration (UF), NanoTrap (NT), and membrane adsorption (MEM) for their suitability to concentrate a wide variety of viral taxa from raw wastewater for PCR detection and sequencing-based metagenomic readouts. We found that to capture a taxonomically diverse virome from wastewater, PEG and VDC outperform all other methods tested in recovery rates, reproducibility, species detection, and captured nucleotide diversity. We observed that different methods exhibit variable concentration efficiencies across taxonomic groups in a reproducible manner, though we could not identify common physiochemical attributes driving this difference. We conclude both PEG and VDC are equally capable at detecting and enriching a broad range of viral taxa, boosting the genomic information potential and reducing blind spots relative to other tested methods. These results advance WBE towards capturing the complex wastewater virome and help guide protocol choices for potential future viral threats.

**Highlights:**

- Virus concentration method choice highly impacts recovered viral communities.
- Polyethylene glycol precipitation (PEG) and vacuum-based direct capture (VDC) detect more viral species with higher nucleotide diversity than other methods tested.
- Enrichment effect of concentration depends on viral taxonomy.
- PEG and VDC yield comparable enrichments of a broad range of viral taxa from wastewater.

## 1. Introduction

Wastewater (WW) is increasingly recognized as a valuable source of information about the health of the contributing population (Choi et al., 2018) including drug use (Huizer et al. 2021), chemical exposure (Vitale, Morales Suárez-Varela, and Picó 2021), biomarkers (Gracia-Lor et al. 2017), and pathogen surveillance (Polo et al. 2020; Kilaru et al. 2023). Of particular interest are viral pathogens, which are shed into the wastewater catchment through feces, urine, and other bodily fluids, and transported to centralized sampling points, typically at local wastewater treatment plants (WWTPs). Such samples can provide an unbiased view of the viruses circulating in a population. Notably, this is not hindered by constraints such as available access and participation in a healthcare system, and symptomatic disease manifestation present in clinical sampling strategies. Wastewater sampling overcomes these limitations by passively sampling the population, assuming the population is connected to centralized wastewater infrastructure. Multiple studies have shown high concurrence between wastewater load and clinical load (Melvin et al. 2021; Amman et al. 2022; Ahmed et al. 2023; Tisza et al. 2023). This has been successfully applied to monitor and track the spread of SARS-CoV-2 variants and outbreaks (Amman et al. 2022, Karthikeyan et al., 2022, Jahn et al., 2022), Mpox (Oghuan et al. 2023; Sherchan et al. 2023), Influenza A (Wolfe et al. 2022), Poliovirus (Tambini et al. 1993), and many other pathogenic targets (Kilaru et al. 2023).

To address the highly diluted nature of viruses in raw wastewater, concentration methods to enrich virus particles are usually applied before nucleic acid extraction. Ideally, a suitable concentration method for a representative surveillance of the circulating virome specifically enriches virus particles regardless of physicochemical characteristics (e.g. size, envelope status, isoelectric point, nucleic acid type, shape) while simultaneously reducing the background of non-viral material and inhibitors. Various approaches have been proposed and are routinely used in wastewater pathogen surveillance programs worldwide (Carcereny et al., 2021; Amman et al. 2022; Wade et al., 2022; Adams et al. 2024; Krogsgaard et al., 2024). These methods differ in their mechanism, viral specificity, ease of use, and scalability. A growing body of literature aims to evaluate the robustness and efficiency of these methods for concentrating virions from wastewater. These studies focus largely on respiratory virus targets like SARS-CoV-2, or surrogates thereof, typically using quantitative PCR as a readout. Previous work comparing viral concentration methods has shown that polyethylene glycol precipitation (Hjelmsø et al. 2017; Barril et al. 2021; Salvo et al. 2021), ultrafiltration (Jafferali et al. 2021; Pino et al. 2021; Farkas et al. 2022; Zheng et al. 2022), and adsorption-elution with an electronegative membrane (Ahmed et al. 2020; LaTurner et al. 2021) yield high recoveries of the measured viral targets. More recent methods such as NanoTrap and vacuum-based direct capture, based on Promega’s VacMan^®^, were also shown to perform well for certain viral targets (Dimitrakopoulos et al. 2022; Girón-Guzmán et al. 2023; Antkiewicz et al. 2024;). Given the variable physicochemical characteristics of virus particles across the viral kingdom, the generalization of these results to newly considered viruses are in question. To date few studies have investigated the effects of concentration methods on the total virome using sequencing based read outs (Hjelmsø et al. 2017; Jiang et al. 2024; Li et al. 2024; Cheshomi et al. 2025). These methods suffer from more delicate molecular biological protocols and are more impacted by inhibitor presence and sample composition. At the same time, metagenomic sequencing allows capture of the full viral diversity, and is already used in WBE applications to track a wide variety of viruses. Assessing the suitability of different concentration methods across this wide range of taxa not only allows evaluation of their suitability for metagenomic applications but can also inform method choices for future emerging viral pathogens.

To evaluate the process of enriching viral nucleic acids at the virome scale, we set out to comparatively analyze five different concentration methods using orthogonal readouts of dPCR against 11 morphologically diverse viral targets, combined with deep sequencing to evaluate overall viromic data quality and identify method-associated biases across a broad spectrum of viral taxa. We then compared the two best in-class methods to confirm the reproducibility of the observed signals across different WWTPs of varying contributing population sizes and different wastewater matrices, offering valuable generalizable insights for future viral WBE applications.

## 2. Materials and Methods

### 2.1 Wastewater Collection

Samples were collected from the inflows of three different municipal WWTPs within Central Europe using activated sludge treatment including nitrification, denitrification and phosphorous removal. Sampling was done volume-proportionally (c.v.v.t – constant volume variable time) using an automatic sampler over a period of 24 hours. The composite samples were subsequently transported to the laboratory where samples were stored at 4°C until further use. All samples were processed within 48 hours.

### 2.2 Experimental Setup

To evaluate the performance of different virus particle concentration methods across the community of viruses in wastewater, we started out with a 24-hour composite wastewater sample from a large, municipal WWTP (serving a population of 2 million), treating the combined sewer effluent of a big city. Supplemental Figure S1 visualizes the experimental setup.

A viral spike-in mix was created consisting of five viruses, selected to include enveloped/non-enveloped, different sizes, genetic nucleic acid types, and shapes by mixing individual stocks of Influenza A (PR-8 strain), Vaccinia (Western reserve strain), LCMV (Clone 13 strain), LCMV (Armstrong strain), Bacteriophage MS2, and PhiX174 at a target concentration of 1×10^5^ genome copies/mL. For details, refer to Table S1. The spike-in mix was added to 600 mL of raw wastewater and equilibrated for 2 h at 4°C with regular inversion every 10 minutes to ensure adequate mixing. An equal volume of spike-in mix was added directly into a 2x volume of DNA/RNA shield for direct TNA extraction.

The spiked wastewater was subjected to five different concentration methods: Membrane adsorption (MEM), ultrafiltration (UF), NanoTrap^®^ (NT), vacuum-based direct capture (VDC), and polyethylene glycol precipitation (PEG). Negative controls of an equal volume of H_2_O (NTC) were run through each concentration protocol to control for lab contaminants and a non-spiked WW sample (NSC) served as a control for background levels of the spike-in viruses already present in the WW. The concentrated wastewater was then used as input for total nucleic acid (TNA) extraction along with three replicates of 500 µL of raw, unconcentrated (UNC) wastewater from both the spiked and non-spiked WW samples. The unconcentrated extracts served as an unprocessed reference, which was expected to be less sensitive due to the reduced input volume, but unaffected by method-specific bias, or sample loss associated with the additional handling steps.

Subsequently, all samples were analyzed for selected viral targets by dPCR and for the bacterial 16S rRNA gene by qPCR (see below) and prepared for sequencing both by untargeted metagenomics and hybrid-capture deep sequencing using the Twist Comprehensive Viral Research Panel (see below).

Finally, we tested the robustness of our findings by expanding our analyses beyond samples of a single large, urban WWTP by including wastewater samples from a second time point and two additional WWTPs in our analysis. To cover different wastewater matrices, we selected a medium-sized, semi-urban WWTP and a small, rural, sewer catchment, serving approximately 55,000 and 10,000 inhabitants respectively, all located within the same geographic region in Central Europe. Samples from these three WWTPs were concentrated with PEG and VDC in triplicates and analyzed by hybrid-capture sequencing,

### 2.3 Concentration Methods

From the spiked wastewater stock, triplicates of 40mL were aliquoted per concentration method, though for NT only 10 mL was aliquoted following suggestions by the manufacturer. After an initial step for pelleting down large solids from the WW, ultrafiltration (UF) was carried out using Amicon Ultra-15 (30 kDa) Centrifuge Filter tubes to concentrate 40 mL to a volume <0.5mL which was directly put into the TNA extraction. For the NanoTrap® (NT) concentration method, virus particles were enriched with Ceres Nanotrap® Microbiome A magnetic particles according to the manual from the manufacturer. MEM samples were processed following a vacuum-filtration protocol with 0.45 µm mixed cellulose ester membrane (Merck Millipore, HAWP04700) as described in Ahmed et al. (2020) with slight modifications. PEG precipitation was conducted as described in Radu et al. (2022) with a few adaptations. Silica membrane adsorption based viral enrichment using a vacuum-based direct capture device (VacMan^®^, Promega) was conducted according to manufacturer’s protocol with a few adjustments. Details on the different concentration methods are given in the supplemental text.

### 2.4 TNA Extraction

TNA extraction was conducted using a magnetic bead based, automatic extraction device (Maxwell RSC; Promega) with the “RSC PureFood GMO and Authentication Kit” (Promega). The cartridges were prepared according to the manufacturer’s instructions. For unconcentrated samples as well as samples enriched using PEG, VDC, UF, and MEM, 500 µL lysis buffer were added to the extraction cartridges. For NT samples, no additional lysis buffer was added. Samples were eluted in 100 µL elution buffer. PEG and VDC samples from the repetition experiment were eluted in 60 µL.

### 2.5 PCR Assays

Quantification of five spiked and six endogenous viruses was performed by dPCR on the QIAcuity dPCR system (QIAGEN) with 26k partition nanoplates. Bacterial 16S rRNA gene (V3-V4 region) quantification was conducted using qPCR on the Applied Biosystems^TM^ QuantStudio^TM^ 6 Pro Real-Time PCR (Thermo Fisher Scientific). Details on the dPCR targets and their viral characteristics can be found in Table S1 and information on cycling conditions, primer and probe information and on analysis in the supplemental text as well as in Suppl. Table S2.

### 2.6 Library Preparation and Hybrid-Capture

15 µL of each extract was used for input to the library preparation. Reverse transcription was carried out with New England Biolabs (NEB) Protoscript II (NEB, E6560L) and NEB Second Strand Synthesis Module (NEB, E6111L) following manufacturer’s protocol. Each cDNA product was equalized to 100 ng as input to the NEBNext Ultra II DNA FS Library Prep (NEB, E7805L) protocol and subsequent hybrid-capture with the Twist Comprehensive Viral Research Panel (Twist Bioscience). Full details can be found in the supplemental text. Both hybrid-captured and untargeted shotgun libraries were sequenced with a paired-end, 200 cycle NovaSeq SP flow cell at the Biomedical Sequencing Facility of the Research Center for Molecular Medicine of the Austrian Academy of Sciences (CeMM).

### 2.7 Data and Code Availability

Sequencing results were processed with an in-house pipeline adapted from Tisza et al., 2023. Full details concerning the workflow and subsequent analyses are found in the supplemental text. All code used in the analysis and figure production can be viewed on Github (https://github.com/thorntonmr/comparative_ww_virome_analysis_manuscript). The data for this study have been deposited in the European Nucleotide Archive (ENA) at EMBL-EBI under accession number PRJEB86516 (https://www.ebi.ac.uk/ena/browser/view/PRJEB86516).

## 3. Results

### 3.1 Quantification of selected viruses and bacterial background by dPCR/qPCR

A comparison of the viral quantifications measured by dPCR for six endogenous and five spiked viruses highlights the pronounced differences between extracts from different virus concentration methods (Fig. 1A, Suppl. Fig. S2). We calculated the recovery performance for all targets by comparing the measured quantity against a theoretical value obtained by multiplying the unconcentrated quantification value by the volume concentration factor (except for rotavirus due to being below the limit of detection in UNC extracts). This number reflects the expected viral copy number in a perfect concentration with no loss relative to the unconcentrated value. VDC had the highest mean recovery of 18.3% followed by NT and PEG (with 12.8% and 9.7%). Overall, most recoveries were less than 20% of the expected value (Fig. 1A). Performances varied between methods with VDC, PEG, and NT outperforming UF and MEM for most targets except for Vaccinia, where NT and MEM showed better performance compared to the other enrichment methods (Fig. 1A, Suppl. Fig. S2, Suppl. Table S1). As targets covered a large range of concentrations spanning over five orders of magnitude, we also investigated the effect of target concentration on recovery. We found that recoveries were largely unaffected by concentration (Suppl. Fig. S3A). When comparing reproducibility between technical replicates, all methods produced comparable coefficients of variation with PEG exhibiting the most uniform variation across all the viruses (Suppl. Fig. S3B).

**Figure 1:**
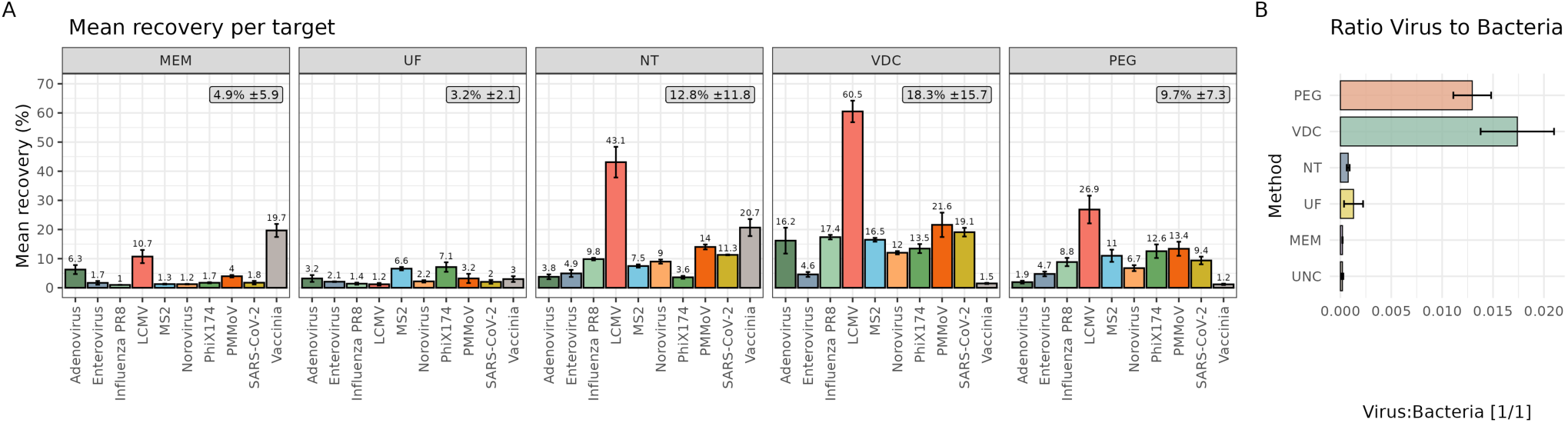
Quantification of target viruses by dPCR across different enrichment methods. Analysis of target virus quantification from wastewater extracts by dPCR assays designed against spiked-in and endogenous viruses. **(A)** Recovery of ten target viruses in extracts from different concentration methods expressed as mean recovery ±standard deviation (error bars) across n=3 replicates. Box information gives the overall average recoveries per method. **(B)** Display of the ratio between sum of all virus target quantifications and 16S rRNA gene content, as a proxy for concentration efficiency of respective methods for virus particles over bacterial cells.

To estimate bacterial contamination, we used 16S rRNA gene quantifications (qPCR). PEG and VDC showed a reduced load of bacterial sequences relative to measured virus suggesting a higher specificity for concentrating viral particles (Fig. 1B).

### 3.2 Virome Assessment by NGS

Hybrid-capture deep sequencing was used to investigate the impact of method selection on the virome detected in wastewater. The initial dPCR quantifications correlated with the relative abundance measures from the dPCR-tested viruses and those included in the hybrid capture panel for sequencing (R² = 0.47-0.84 for different methods; Suppl. Fig. S5).

All methods had highly reproducibly species resolution data with PEG and VDC standing out in the number of species detected in the low abundance range (Fig. 2A). Analyzing the overall picture of sample compositions at superkingdom level revealed that all concentration methods increased the number of viral reads compared to the unconcentrated samples. PEG had the highest percentage of viral reads followed by VDC (28.9%±3.73 and 23.8%±0.83), as well as the lowest percentage of reads assigned to bacterial taxa (28.6%±1.75 and 41%±1.3). UF, MEM, and NT all had roughly half the percentage of reads on target (12%, 14.2%, 15.4%) and bacterial content load followed the trend observed in the 16S rRNA gene dPCR, suggesting the 16S rRNA gene quantification gives a reliable indicator of background sequence contamination (Fig. 2B).

**Figure 2.**
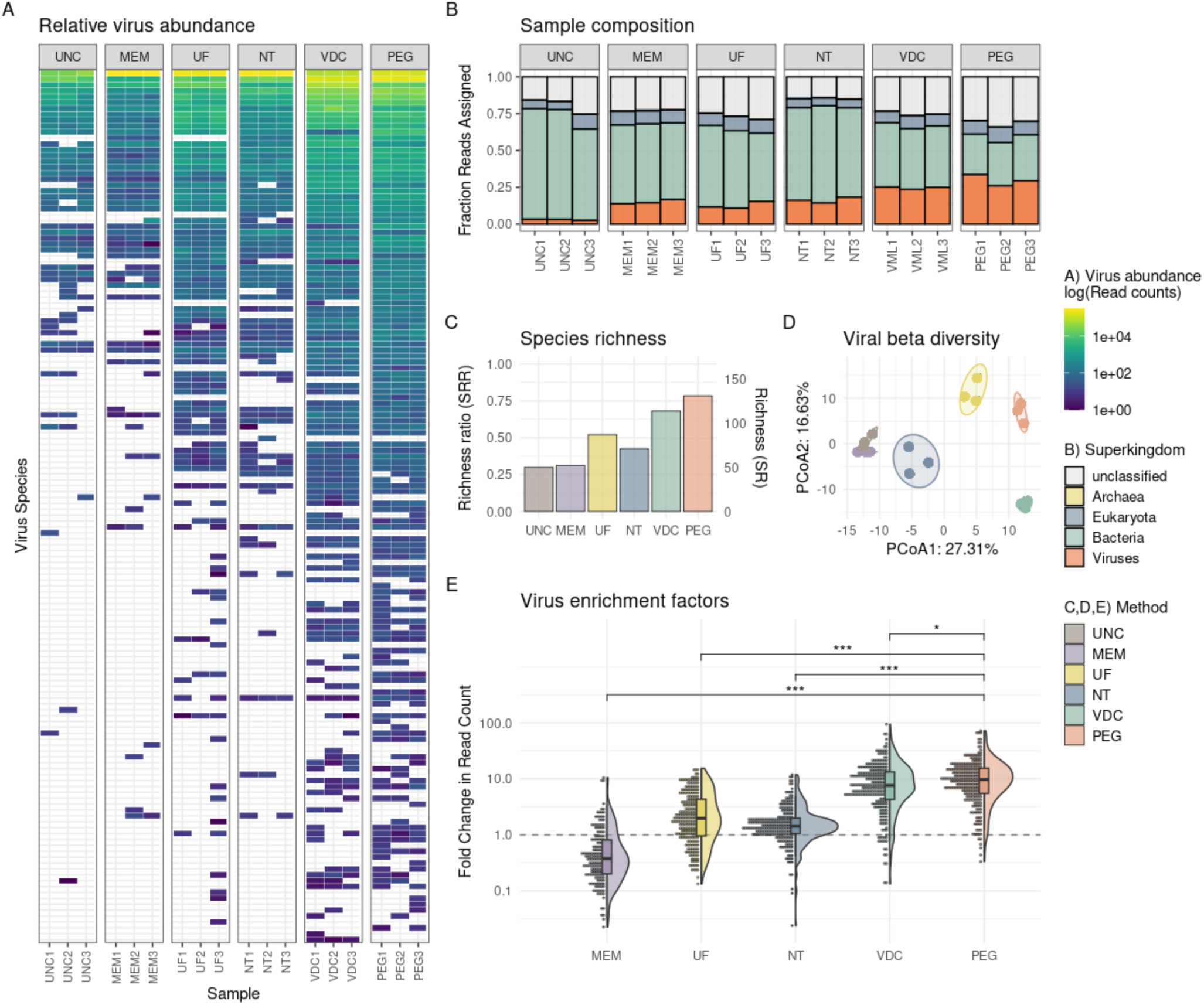
Sequencing results of enriched viral libraries. **(A)** Heatmap showing viral species abundance with tile color indicating the log read count. Species names were removed for visualization purposes but are shown in Suppl. Fig. S6. **(B)** Composition of each sample by superkingdom via read assignment by Kraken2. **(C)** Species richness and richness ratio calculated using Chao’s lower bound estimate. **(D)** Principal component analysis of the Aitchison distance for each method. Points are replicates of the same method. **(E)** Enrichment factor (read count ratio) of each method relative to the mean unconcentrated read count. Significance bars are only shown for comparisons to PEG for visualization purposes. Pairwise significance was tested with a Kruskal-Wallis rank-sum test and post hoc Dunn’s test with the Holm-Bonferroni correction for multiple testing. Significance codes are as follows: * p<0.05, *** p<0.001.

Expected species richness (SR) was calculated across the full dataset (all methods, all replicates) with Chao’s lower bound estimator. We then compared the observed species count per method to this overall estimate to obtain the species richness ratio (SRR). PEG recovered 76% of the maximum detected species number, followed by 67% for VDC and 51% for UF. All other methods detected fewer than 50% of the theoretically detectable viruses (Fig. 2C).

Given that the concentration methods differ in their mechanism of action, the resulting viral community recovered by each method may differ in both composition and abundances of its individual taxa. To assess this, beta diversity measures were calculated for the viral communities of each method and visualized with a principal coordinate plot, showing that the replicates per method separate in distinct, tight clusters, except for MEM and UNC which cluster together (Fig. 2D). This supports the notion that the evaluated concentration methods reproducibly produce different outcomes for different taxa.

To quantify the enrichment effect of each method on detected viral taxa, we used the mean read count per viral strain detected in the three unconcentrated samples as a reference point. Ratios of the read counts per concentration method were compared to the unconcentrated mean to obtain an enrichment factor. PEG outperformed all methods, followed closely by VDC, in enriching read counts per virus. MEM often returned lower read counts than the unconcentrated samples, suggesting a major reduction in the data quality of many viral taxa (Fig. 2E).

### 3.3 Assessing taxonomic or physical attribute biases

It would be highly beneficial to identify specific biases associated with a particular method to unveil the limitations of each method. To this end, we sought to identify taxonomic or physical attribute biases associated with method performance. We observed that the method performance was highly impacted by taxonomy. PEG was able to significantly improve enrichment in all but Papillomaviridae and was either significantly better or not significantly worse than all methods for all taxa outside of Adenoviridae where VDC performed best (Fig. 3).

**Figure 3.**
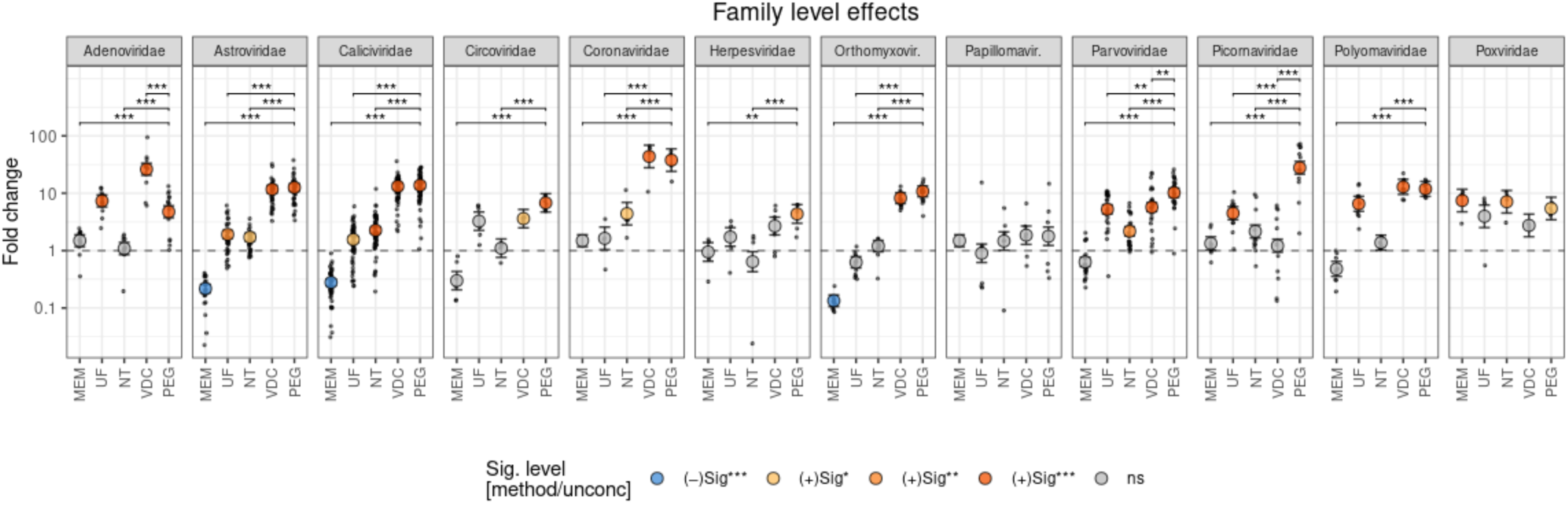
Effects of concentration method on viral enrichment and diversity. Modelling results of enrichment factors with points representing model estimates, point color indicating a significant enrichment compared to the unconcentrated values, and significance brackets showing significant difference in enrichment between methods within a family. Within-family significance is shown only relative to PEG (for visualization purposes). Jitter points show the raw ratio of method to reference read count per strain detected and error bars show the 95% confidence interval. Significance codes are as follows: * p<0.05, ** p<0.01, *** p<0.001.

While taxonomy should implicitly capture the morphological variations between viruses, we also directly investigated the effects of the physical attributes of each virus collected at the genus level to understand if there were sizable biases associated with a particular method. PEG, in addition to the overall highest enrichment, showed a very stable performance across all characteristics, only differing significantly in its improved enrichment of RNA viruses over DNA viruses. VDC was slightly more variable with an improved enrichment in medium sized particles and decreased performance with large particles (Suppl. Fig. S7).

### 3.4 Investigating captured nucleotide diversity

For evolutionary and epidemiological applications of deep wastewater sequencing it is often important to understand not only how many viral taxa are identified, but also how much diversity can be captured by each method for each taxon. An effective concentration method should capture and concentrate more viral particles per taxa, increasing the chances to capture more representative nucleotide diversity within a particular viral sequence. Calculation and pairwise comparison of the diversity metric π (as defined in Nelson & Hughes, 2015 equation 4) revealed that PEG significantly increased captured diversity over all methods except for VDC, where no significant difference was observed (Fig. 4).

**Figure 4:**
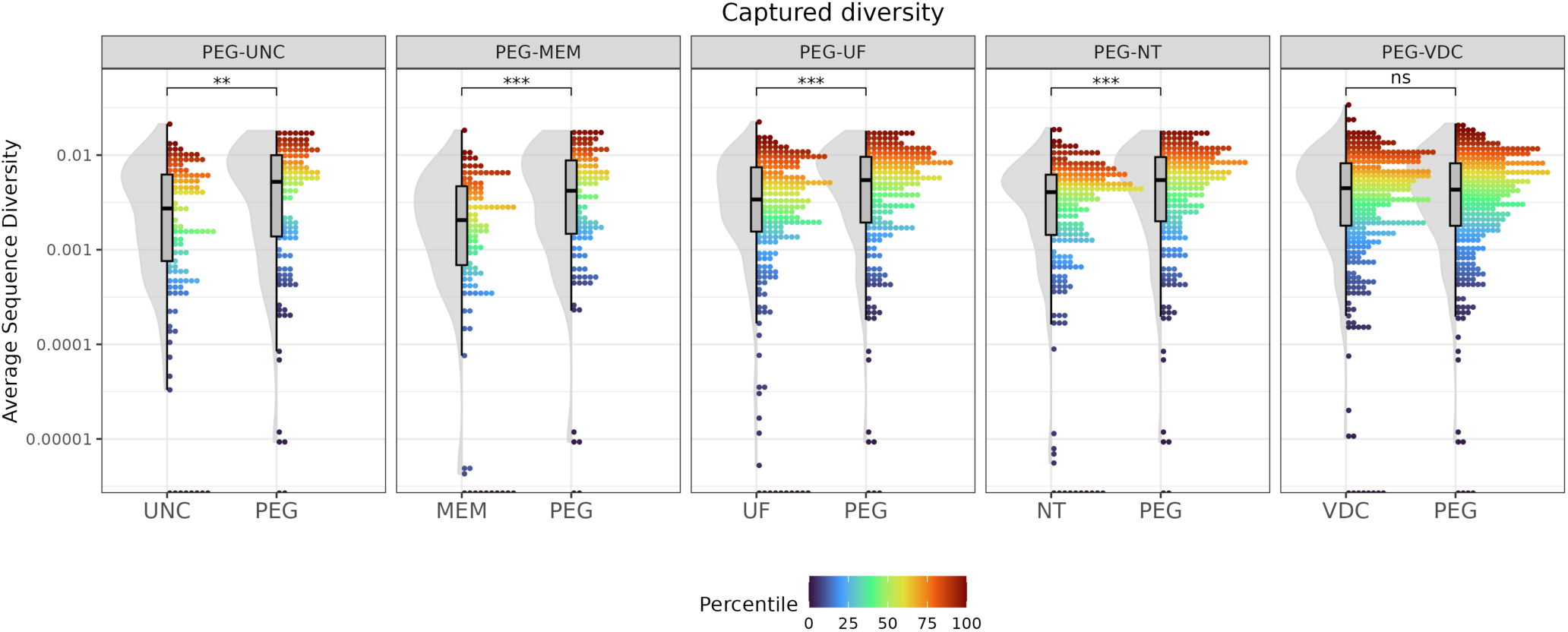
Captured nucleotide diversity. Average per sequence nucleotide diversity of each method compared to PEG. Significance was determined with the Kruskal-Wallis rank-sum test. Dots indicate a single reference sequence and color is scaled by within-sample percentile. Significance codes are as follows: * p<0.05, **p<0.01, ***p<0.001.

### 3.5 Comparing Best-in-class Methods

Finally, we evaluated how reproducible the trends we observed were across different WWTPs and time points. We observed a reproducible method bias across WWTPs where viruses that tended to yield more reads (considered better enriched) for PEG or VDC did so repeatedly across varying WW matrices. The lowest correlation was at the strain level (R=0.53), which can be expected given the increase in stochastic effects due to read assignments and presence at strain level but increases to 0.68 at the family level indicating that this trend holds across repeat measurements (Fig. 5A-D).

**Figure 5.**
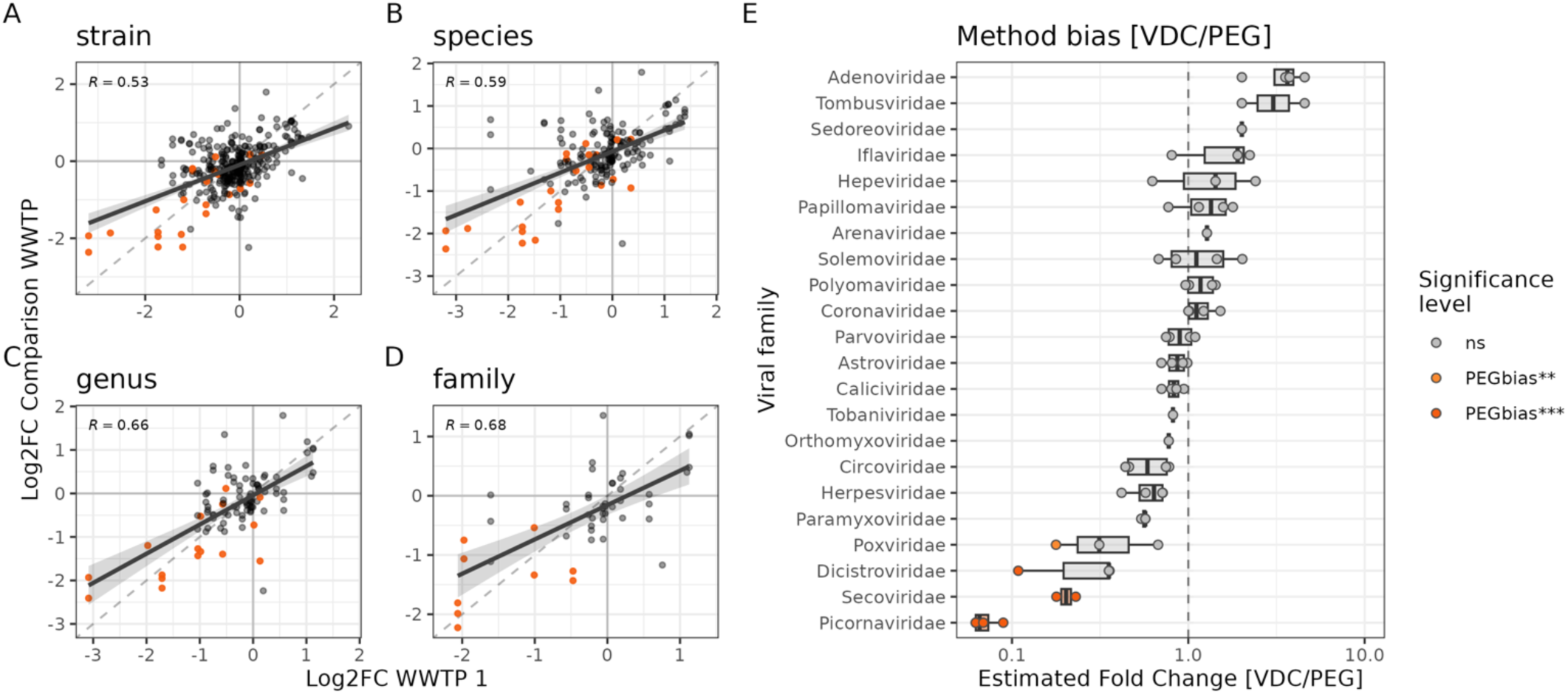
Comparison of PEG and VDC across multiple WWTPs. **(A-D)** Reproducibility of method bias across WWTPs. The x-axis values are log_2_ read ratios [VDC:PEG] in the original experiment and the y-axis values are log_2_ read ratios for all three additional WWTPs tested. Point color indicates strains within each significantly biased family from the model output in panel E and the dashed line represents the perfect correlation. Correlation coefficients were computed from a linear regression through the log_2_ transformed dataset. **(E)** Marginal effects of modelled method bias [VDC:PEG] at the family level in each WWTP. Points represent a bias estimate at a single WWTP and color represents the significance of the bias (no significant VDC biases were observed).

Estimates of method bias from a negative binomial model fit on the read counts per viral strain for each method were reproducible across WWTPs with little difference in the methods performance at the family level. Notably, only taxa within Picornaviridae were significantly enriched in PEG with more than 10-fold more reads assigned. Adenoviridae sub-taxa tended to be more enriched in VDC in all WWTPs. However, after controlling for multiple testing, no significance was confidently assigned to this interaction (Fig. 5E).

## 4. Discussion

This study aimed to expand upon previous comparisons of wastewater concentration methods to optimize the accessible information potential for comprehensive wastewater viromics. dPCR can provide a more direct measurement of concentration efficiency, and is a fast, cheap, and sensitive method for quantifying viral nucleic acids. However, it is limited by prior sequence knowledge, sensitivity to mutation, and the number of targets that can be measured in a single run. Our dPCR data showed that VDC performed best in overall recovery, trailed by NT and PEG. Support for VDC’s high performance in dPCR data is shown in an independent study showing that this method outperformed PEG, UF and MEM in SARS-CoV-2 recovery (Dimitrakopoulos et al. 2022).

However, it is often important to cast a much wider net when profiling viruses from wastewater to identify and investigate multiple taxa, including taxa for which little *a priori* knowledge is available, or when targeted methods are hampered by sequence variability within the target species. Virome-scale sequencing provides the advantage over dPCR in such cases allowing for both detection and probing of the genomic information across all detectable taxa present in a sample. Given the compositional nature of sequencing data, the detection success and accompanying depth of genomic information are heavily affected by the specificity of the method for viral particle enrichment. Method choice highly impacts the ability to detect viral taxa as exemplified in Figure 2C and Supplemental Figure S9, which indicate the level of false negatives an analyst may encounter when using a sub-optimal method. While dPCR metrics provided a rough predictor of sequencing performance, it was necessary to consider the 16S rRNA gene measurement to account for background sequence contamination. We observed large differences in the bacterial content per method suggesting variability in the virus specificity. This greatly hindered the sensitivity for detecting viral sequences. In the case of NT, viral recoveries alone were poor indicators of the downstream sequencing performance (Fig. 2C).

Overall, we obtained several lines of evidence indicating that PEG and VDC are effective methods for concentrating a diverse set of viruses independent of physiochemical properties. PEG, VDC, and UF methods all had an initial centrifugation step to remove large particles from the sample while MEM and NT both used direct inputs without the initial centrifugation. PEG showed the highest viral specificity, which may be attributed to bacterial removal during the initial centrifugation step (Hjelmsø et al. 2017). The exact mechanism of action for PEG remains poorly characterized though the prevailing hypothesis is that virus particles are aggregated in the extra-polymer space and precipitated out after exceeding their solubility (Ingham 1990). In support of our findings, Yamamoto et al. (1970) found that PEG precipitation efficiency is independent of viral concentration, pH, and ionic strength and Atha and Ingham (1981) showed that PEG has very little interaction with proteins supporting a lack of discriminatory features where performance bias can arise.

VDC is essentially a large-scale extraction and purification, so there is not an obvious avenue for large bias introduction across taxonomy or physical particle attributes. Given this, it is surprising to observe such a high virus to bacteria ratio though likely the initial centrifugation influenced the observed viral specificity.

UF relies solely on size exclusion by allowing liquids to pass through the 30kDa filter but retaining everything else and reducing the overall volume. Poor overall performance from UF could be attributed to viral adsorption onto the membrane, which may also leave room for the preferential adsorption of certain particle types. Co-concentration of inhibitors present in the samples could also affect the assays used to produce the quantifications and sequencing results, and though inhibitor load was not explicitly investigated here, the enrichment rate for RNA viruses was significantly less than that of DNA viruses in UF and reverse transcription is a step sensitive to the presence of inhibitors (Suppl. Fig. S7).

MEM relies on the charge properties of viruses requiring a negatively charged particle to bind via MgCl_2_ mediated salt-bridging to the electronegative membrane (Cashdollar & Wymer, 2013), which can be impacted by properties of the wastewater matrix (pH, turbidity). The limited performance of MEM relative to the other methods found in the present study is supported by dPCR-based results from Antkiewicz et al. (2024) which showed VDC, NT, and PEG outperform MEM methods when quantifying SARS-CoV-2 and bovine coronavirus.

NT relies on chemical affinity capture of viral particles and has been shown to vary in capture rate based on virus size and envelope composition (Lin et al. 2020). NT has also been shown previously to detect fewer species than PEG following hybrid-capture sequencing (Jiang et al. 2024). While our data suggests NT to reliably quantify a range of dPCR targets, it falls short in the sequencing results. The NT Microbiome A Particles used in this study enrich viral particles as well as certain bacteria expanding the range of targets for concentration, at the expense of selectivity which can be beneficial or detrimental depending on the desired use-case. A benefit to NT is the shorter hands-on time and ease of automation allowing this method to be readily scaled.

We could not identify systematic differences between the two top method’s performances outside of the enrichment of Picornaviridae in PEG. This viral family contains enteroviruses which are of particular relevance, given that many WBE programs screen for poliovirus sequences (Brouwer et al., 2018; Whitehouse et al., 2024). However, the PEG bias seems to be driven by a subset of genera within Picornaviridae, namely *kobuvirus* and *salivirus*, while enterovirus bias is largely unaffected by method choice (Suppl. Fig. S8).

This study has several limitations. We employed protocols that were previously shown to perform well in published literature. However, it is likely that modifications to these protocols could boost or hinder performance. For instance, Farkas et al. (2022) showed that addition of beef extract-sodium nitrate elution to PEG precipitation greatly improved viral recovery. Even though we followed protocols for MEM similar to those used in Ahmed et al. (2020) and LaTurner et al. (2021), our findings contradict the high levels of recovery observed in MEM and low recoveries from PEG. Furthermore, there are additional methods to concentrate viruses which were not included as it was out of scope of this study to include all potential methods and variations.

Linking the observed biases to physicochemical properties rather than the taxonomic identity of detected viruses would allow more generalizable results as the physical characteristics of the virus directly interact with the WW matrix and concentration method’s mechanism of action. However, due to inherent class imbalances these properties are difficult to adequately model leaving taxonomy as a more balanced alternative. In general, we did not observe much variation on the genus level though this observation is only meaningful if a representative portion of the genus was captured in our sample set. Despite this limitation, PEG and VDC were not outperformed by any method across all observed taxonomic groups supporting the notion that either method could be the initial choice for profiling novel viruses, unless alternative approaches are systematically evaluated and demonstrated to be superior.

## 5. Conclusion

In summary, we investigated the ability of five widely used WW concentration methods to effectively concentrate a broad range of viruses. dPCR was used to assess raw quantifications and recoveries of each method, while deep hybrid-capture sequencing was used to examine performance across the detectable human virome in WW. We found PEG and VDC to be the best-in-class methods for viral concentration providing high relative recoveries and technical stability across a broad range of viral taxa, and the highest capture of nucleotide diversity within detected viral sequences. We observed reproducible trends with little systematic differences between PEG and VDC across independent WWTP sampling, highlighting both methods to be viable choices for virome-scale concentration in support of WBE.

## Acknowledgements

We would like to thank Professor Regina Sommer for kindly providing the PhiX174 and MS2 phage stocks, the Biomedical Sequencing Facility of the Research Center for Molecular Medicine for the sequencing of all libraries in this study, Helene Loges for conducting the 16S rRNA gene quantifications, and the WWTP operators for providing samples. The work was financially supported by the supported by the Austrian Science Fund, Cluster of Excellence COE7 and the FFG KIRAS project 909310 granted to AB.

## Supplemental Text

### Concentration Methods

#### - Ultrafiltration (UF)

40 mL of WW was spun down for 10 minutes at 3,000 × *g* to pellet large solids that would clog the membrane. Amicon Ultra-15 (30 kDa) Centrifuge Filter tubes (Merck Millipore, UFC903024) were flushed with 10 mL of H_2_O and spun down for 10 minutes at 4,750 × *g* prior to loading samples. 15 mL of each sample was then loaded into a centrifuge tube and spun down in increments of 10 minutes at 4750 × *g* until >1 mL of concentrate remained. If the filter clogged, the tubes were inverted a few times to break up the solids collection. The flow through was discarded and another round of volume added until the full 40 mL was concentrated to a volume <0.5 mL which was directly input to TNA extraction.

#### - NanoTrap^®^ (NT)

100 µL of Ceres NanoTrap^®^ enhancer reagent 2 was added to 10 mL of wastewater and vortexted for several seconds. Subsequently, 150 µL of Ceres Nanotrap® Microbiome A particles was added and mixed by inverting 2-3 times. Samples were incubated at room temperature for 10 minutes with an additional inverting step after 5 minutes. The tubes were put in a magnetic stand for 3-5 minutes allowing the magnetic particles to aggregate along the wall of the tube. The supernatant was discarded before the tubes were removed from the magnetic stand and 700 µL of lysis buffer was added. After mixing the particles with the lysis buffer using a pipette, the tubes were put back into the magnetic stand for particle separation. The remaining buffer containing the viral material was loaded into the cartridge for extraction.

#### - Membrane-Adsorption (MEM)

MEM samples were processed following a protocol described in Ahmed et al. (2020) with slight modifications. Briefly, 25 mM MgCl_2_ was added to each sample 30 minutes prior to concentration and inverted to mix. Filter funnels, housing, and forceps were sterilized with a propane torch prior to loading the samples. A 47 mm, 0.45 µm mixed cellulose ester membrane (Merck Millipore, HAWP04700) was added to the sterilized vacuum filter funnel and the full 40 mL sample volume was run through each membrane. Once the full volume had passed through, the membranes were cut into small pieces with sterilized scissors and placed into bead-beating tubes with 500 µL of lysis buffer (from TNA extraction kit, see 2.4 in main text). Samples were run on the FastPrep-24 homogenizer (MP Biomedicals LLC) using three cycles of 20 s at 6 m/s, then spun down for 10 minutes at 10,000 x *g* and retained lysis buffer used for TNA extraction.

#### - PEG precipitation (PEG)

PEG precipitation was conducted as described in Radu et al. (2022) with a few adaptations. In brief, 47 mL of well mixed wastewater was centrifuged at 4,500 × g for 30 minutes at 4 °C. 40 mL of the supernatant was subsequently mixed with 4 g PEG and 0.9 g NaCl By shaking until chemicals were fully dissolved before centrifugation at 12,000 × g at 4 °C for 1.5 h. The supernatant was decanted and centrifuged for 5 minutes at 12,000 × *g* at 4 °C. Residual liquid was removed, the remaining pellet resuspended in 500 µL of a CTAB buffer - H_2_O mix (4:3) and transferred to a 1.5 mL reaction tube. After another centrifugation for 5 minutes. at 16,000 × *g*, the supernatant was loaded into the extraction cartridge.

#### - Vacuum direct capture (VDC)

Silica membrane adsorption based viral enrichment using a vacuum-based direct capture device (VacMan^®^, Promega) was conducted according to manufacturer’s protocol with a few adjustments. In brief, 0.5 mL of Protease Solution was added to 40 mL of wastewater sample, mixed by inverting and incubated at room temperature for 30 minutes. Samples were subsequently centrifuged at 3,000 × *g* for 10 minutes for solid separation. The supernatant was divided into two 50 mL tubes. To each of the tubes containing 20 mL sample, 6 mL Binding Buffer 1, and 0.5 mL Binding Buffer 2 was added. Post inversion, 24 mL of isopropanol was added and again mixed by inversion. PureYield binding columns were added to the VacMan device and the mixture of the two 50mL tubes added gradually. Vacuum was applied until the whole volume passed through the column. Subsequently, 5 mL of Column Wash 1 was vacuumed through the column, followed by 20 mL of Column Wash 2. The vacuum was turned on for additional 30 s to pass the remaining solution. The elution device was assembled and connected to the column before twice 250 µL of Nuclease-Free water (preheated to 60 °C) was added to the column and vacuumed through the column for 1 minute each. The whole pre-extract which was collected in a 1.5 mL reaction tube was added to the extraction cartridge.

### 16S Quantifications

Reaction mix was prepared by combining 5 µL of Master Mix, 0.5 µL of each primer (10 µM working solution; Forward: S-D-Bact-0341-a-S-17; Reverse: S-D-Bact-0518-a-A-17) and 2 µL H_2_O (molecular grade) with 2 µL DNA template/standard/negative control (molecular grade H_2_O) for 10 µL total volume approaches. All samples/standards/negative controls were run in technical duplicates under the following cycling conditions: 10 minutes at 95°C followed by 45 cycles of alternating 15 s at 95 °C and 1 minute at 60 °C. Subsequently, a melting curve analysis was conducted for additional quality control. A standard curve was prepared by diluting a purified linearized recombinant plasmid containing the 16S rRNA gene target (adapted from plasmid pBELX described in Bellanger et al., 2014; produced by Xavier Bellanger & Christophe Merlin) to concentrations reaching from 10^8^ to 1 copies/well. Highest standard dilutions (1 and 10 copies/well) were excluded for standard curve calculations. Amplification efficiency was between 90% and 110% and R^2^ between 0.99 and 1.00. All samples were measured in 1:10 and 1:100 dilutions (with molecular grade H_2_O) to check for inhibition. Mean Ct-values obtained from technical replicates of 1:10 diluted samples were subtracted from means of 1:100 diluted samples. These values were further subtracted from the value that would ideally correspond to a 10-fold dilution (LOG(10)/LOG(2) = 3.322). In case the resulting factor showed a value >1, the measurement was considered as inhibited and the 1:100 dilution was used for further calculations. On the other hand, if the factor showed a value ≤ 1 or the 1:100 result showed values below LOQ, the 1:10 dilution value was used. LOQ was defined by the lowest standard concentration (10^2^ copies/well) used for the standard curve calculation for the respective qPCR run.

### Library Preparation and Hybrid Capture

15 µL of each sample was taken forward for sequencing prep. First, strand synthesis was performed using Protoscript II (NEB) with 5 µL random hexamers (60 µM) and incubation at 25 °C for 5 minutes, 42 °C for 1 hour, and inactivation at 80 °C for 5 minutes. Second strand synthesis was performed for 1 hour at 16 °C using NEB Second Strand Synthesis Module with the heated lid off. Following reverse transcription all samples were cleaned up with 1.0X volume SPRI beads and quantified with a Qubit dsDNA High Sensitivity Assay (Invitrogen). Library prep was performed following the protocol for NEBNext Ultra II DNA FS Library Prep. Inputs were standardized to 100ng and all volumes adjusted to 26 µL. Fragmentation was performed for 20 minutes at 37 °C, followed by 30 minutes at 65 °C. Barcode addition was performed with 7 indexing PCR cycles and PCR products were cleaned up with 0.9x SPRI beads. Equimolar volumes of each library were pooled and set aside for untargeted shotgun metagenomics. Hybrid-capture library pools were prepared by pooling 100ng of each sample in two pools (200 ng total per sample) to reduce the batch effects of hybrid-capture with different sample mixes. Hybrid-capture was performed following the Twist Target Enrichment Standard Hybridization v2 protocol with 14 post-capture PCR cycles. All libraries were sequenced with a paired end, 200 cycle NovaSeq SP flow cell at the Biomedical Sequencing Facility of the Research Center for Molecular Medicine of the Austrian Academy of Sciences (CeMM).

### Bioinformatic Analysis

To ensure comparability across methods, the resulting sequencing libraries were down-sampled to match the smallest library size, accounting for the effect of sequencing depth on species detection, particularly in the low-abundance range (Suppl. Fig. S4). Downsampled reads were quality controlled, trimmed, and deduplicated using fastp (Chen et al., 2018). To remove common contaminant sequences trimmed reads were then mapped to a set of human (GRCh38.p14) and NCBI UniVec references using Bowtie2 (Langmead & Salzberg, 2012) and reads mapping to the contaminant references were excluded from downstream analysis. Clean reads were then processed with an in-house pipeline adapted from Tisza et al., 2023. Briefly, reads were aligned with a 2-pass mapping approach using a database constructed specifically to represent taxa contained in the Twist Comprehensive Viral Research Panel. Reads were mapped to the database using minimap2 (Li, 2018) allowing a maximum 10 secondary alignments at >60% of the primary alignment score. Mappings were filtered with msamtools (https://github.com/arumugamlab/msamtools) to retain reads >60 bp, with an alignment length ≥80%, and nucleotide identity ≥90%. Uniquely aligned reads were indicative for presence of the respective reference sequence. Uniquely aligned reads were defined as reads for which the second-best possible alignment had an edit distance greater than one away from the top alignment. The number of multimapping reads, i.e. non-uniquely aligned reads, was counted per reference sequence as a percentage of total reads mapping to that sequence, as well as percent genome coverage and reference sequence length. The reference set was then filtered per sample to retain only sequences that had ≥20 reads mapped, ≥40% or ≥800 bp of the genome covered by at least one read, and less than 98% multimapping rate. To account for false positives due to short, conserved regions the multimapping threshold was decreased to 80% for sequences longer than 5000 bp with ≤2500 bp covered at 1x requiring more unique reads to be considered a true positive. The initial reference set was reduced to include only sequences passing these detection filters and reads were remapped to the reduced reference set. Mapping reads were then subsampled with seqtk (https://github.com/lh3/seqtk) and used with Salmon (Patro et al., 2017) to estimate per-sequence quantification. A last filtering step was performed with prior thresholds and additionally to remove reads with a Salmon transcript-per-million of zero and final statistics were reported with a custom R script. All subsequent analyses used the probabilistic readjusted read counts from Salmon unless otherwise stated. The analysis pipeline and respective filtering thresholds were evaluated using simulated read data and direct extractions of viral stocks to balance sensitivity and specificity (data not shown). Variants were called using LoFreq (Wilm et al., 2012) with default settings. All statistical analyses were conducted in R (R Core Team, 2024).

### Virome Assessment by NGS

To obtain an overall picture of sample compositions at the superkingdom level, kraken2 was used to assign taxonomy to read fragments using the comprehensive PlusPFP database (Wood et al., 2019) using a read confidence score of 0.3.

### Assessing taxonomic or physical attribute biases

A linear model was fit on log_10_ transformed enrichment factors with method and viral family as interacting predictors. We computed the marginal effects and subsequent hypothesis tests using R package *marginaleffects* (Arel-Bundock et al., 2024). We then compared the method enrichment relative to the unconcentrated reference values and the relative enrichment of each method to the others within each viral family, after controlling for family-wise error rates using the Holm-Bonferroni correction (Holm 1979). We also collected physical attribute information for each viral genus from viralzone.com. We summarized each genus by size classes (<75 nm = small, between 75 and 150 nm = medium, larger than 150 nm = large) using the midpoint if a range was provided, enveloped status, and nucleic acid type and included these as predictors in a separate model of enrichment performance.

### Investigating captured nucleotide diversity

We calculated the diversity metric π (Nelson & Hughes, 2015 equation 4) averaged across all sites covered at >10x depth for each reference sequence detected. To provide a comparable dataset, each pairwise method comparison was conducted on all sequences found in all three replicates in both methods. Using the Kruskal-Wallis rank-sum test, each method was compared only to the top performer in sequencing readouts, e.g., PEG, to reduce the number of tests.

### Comparing best-in-class Methods

A method bias metric was calculated as the ratio between read counts per viral strain in VDC and read counts of the same strain in PEG. This metric was calculated in all three of the newly sampled WWTPs and compared to the values obtained in the initial experiment aggregating the read counts at increasing taxonomic levels. Correlation coefficients were calculated on the log-scale to equally weight points below 1 (PEG-biased) and above 1 (VDC-bias). For the comparison of the five enrichment methods, effects on taxonomy were assessed as in Figure 3. Given that this analysis does not compare enrichment factors to an unconcentrated reference, we applied a model more suitable for overdispersed read counts often found in NGS data (Robinson & Smyth, 2007). To this effect, a negative binomial model with a log link function was fitted on the read counts per viral stain for PEG and VDC with method, viral family, and WWTP as predictors using the R package MASS (Venables & Ripley, 2002).

## Supplemental figures

**Supplemental Figure S1:**
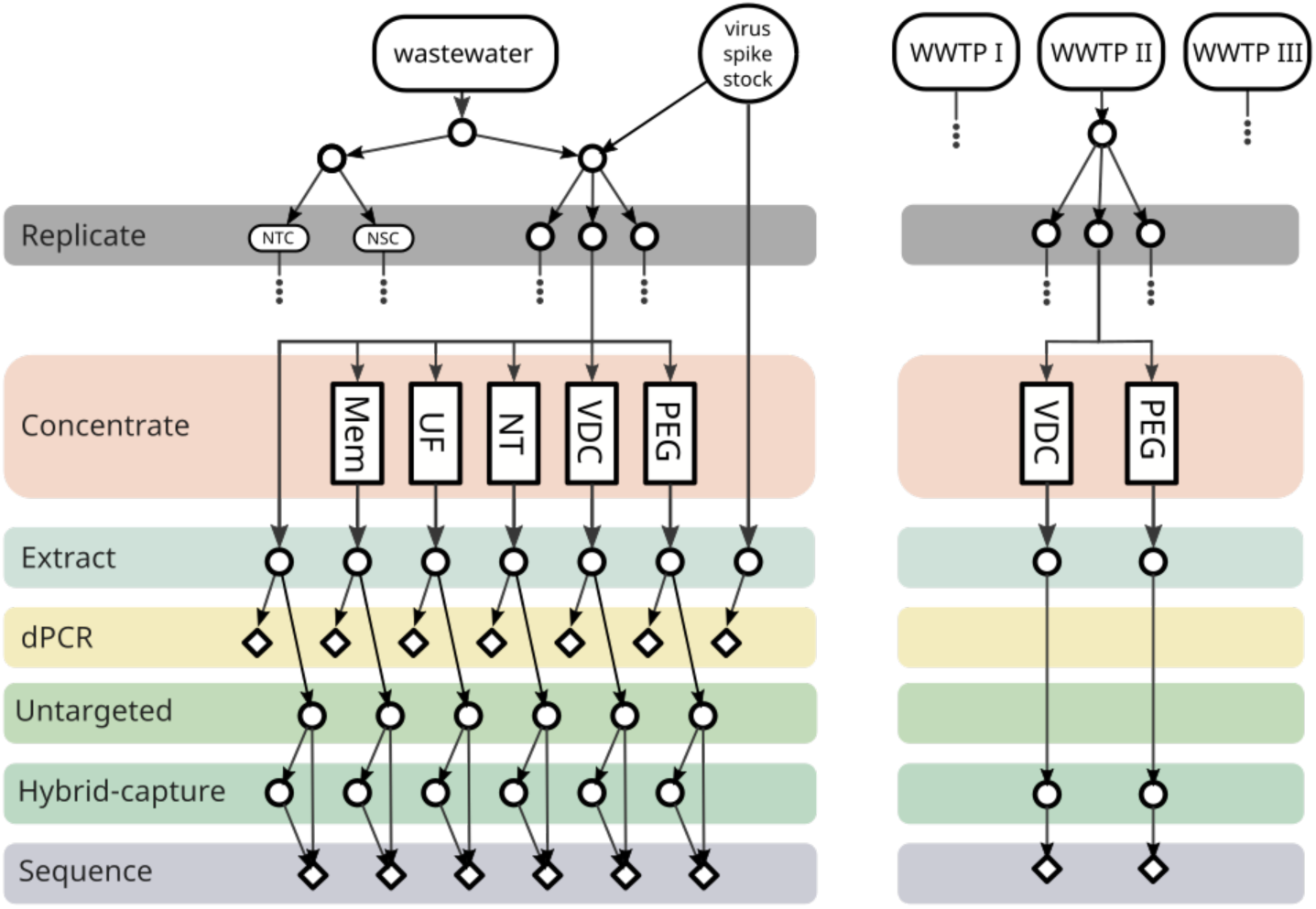
Experimental Design.

**Supplemental Figure S2:**
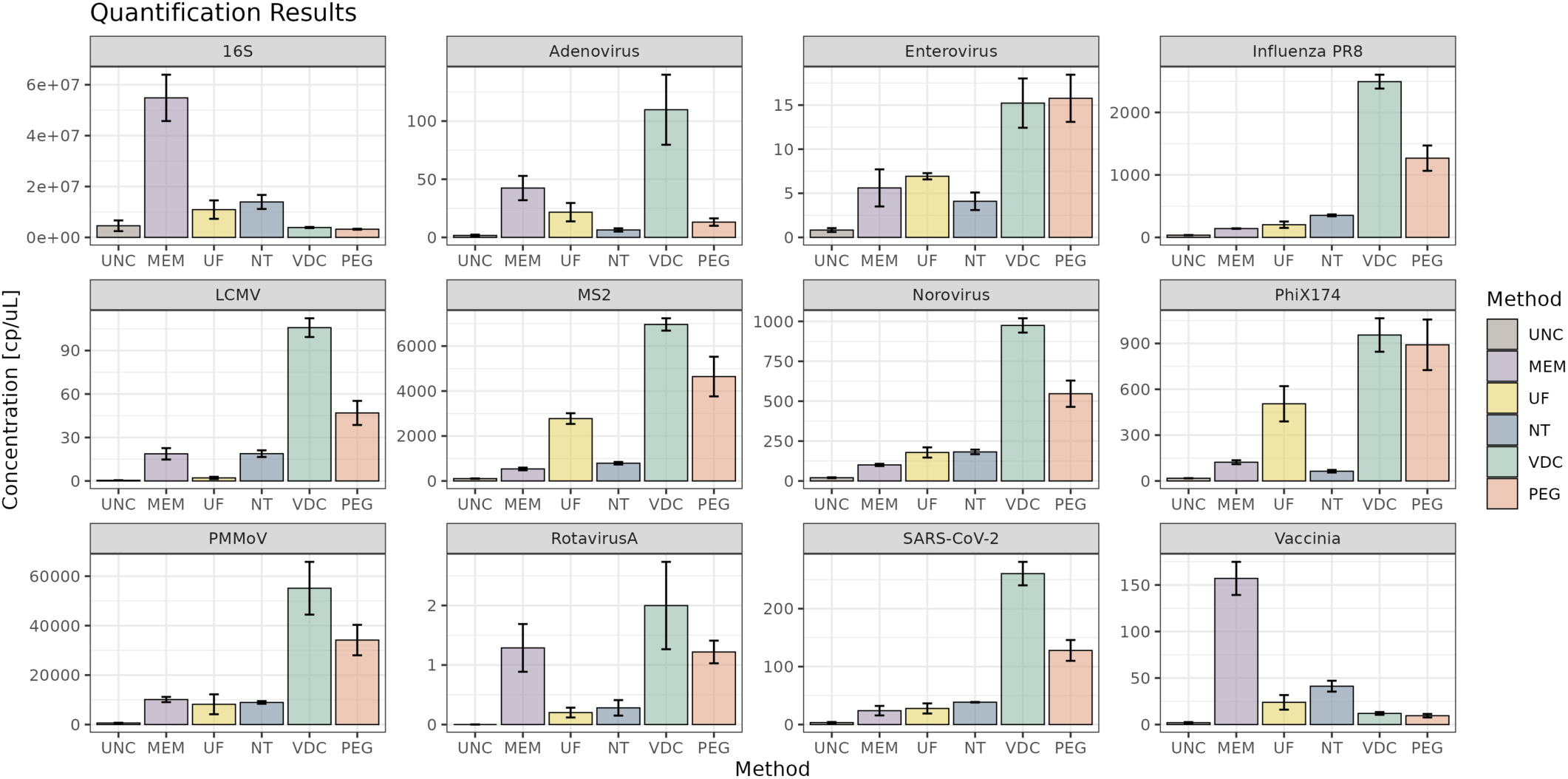
Raw dPCR Quantifications. Quantifications for each measured target are expressed as the mean concentration across all three replicate extracts. Error bars represent the standard deviation. Quantifications are not normalized for input sample volumes here (0.5 mL for UNC, 10 mL for NT, and 40 mL for all other methods).

**Supplemental Figure S3:**
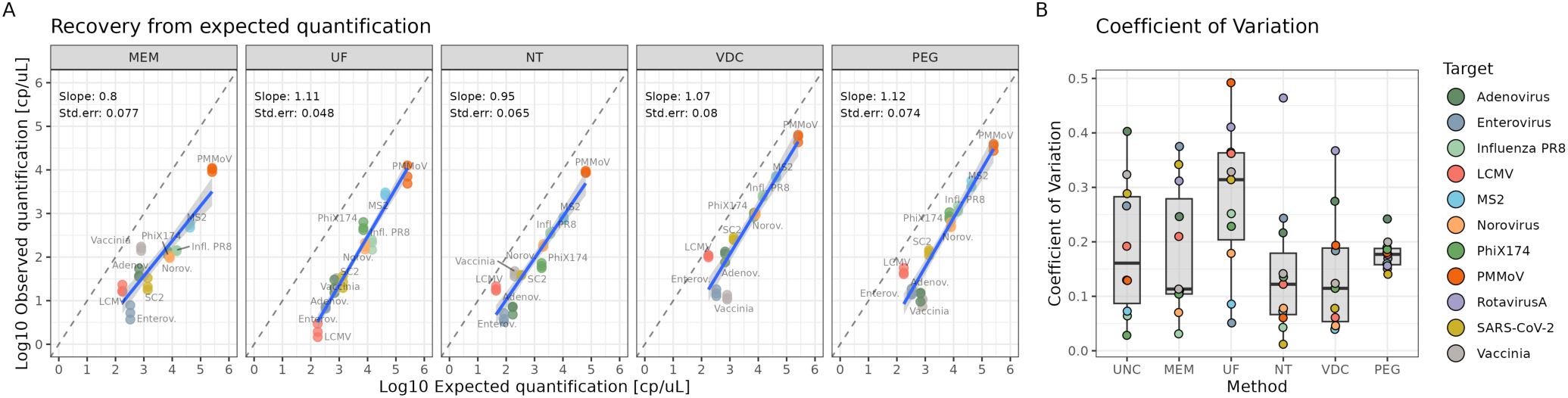
Reproducibility across concentration and technical replicates. **(A)** Scatterplot depicting the estimated recoveries of each target across the range of target concentrations. A linear regression line was fit through log10 transformed concentration values and slope and standard error shown per method. Slopes closer to one indicate lesser impacts of concentration on recovery. **(B)** Coefficients of variation (standard deviation / mean) for each target per method. Colored dots are individual target values.

**Supplemental Figure S4:**
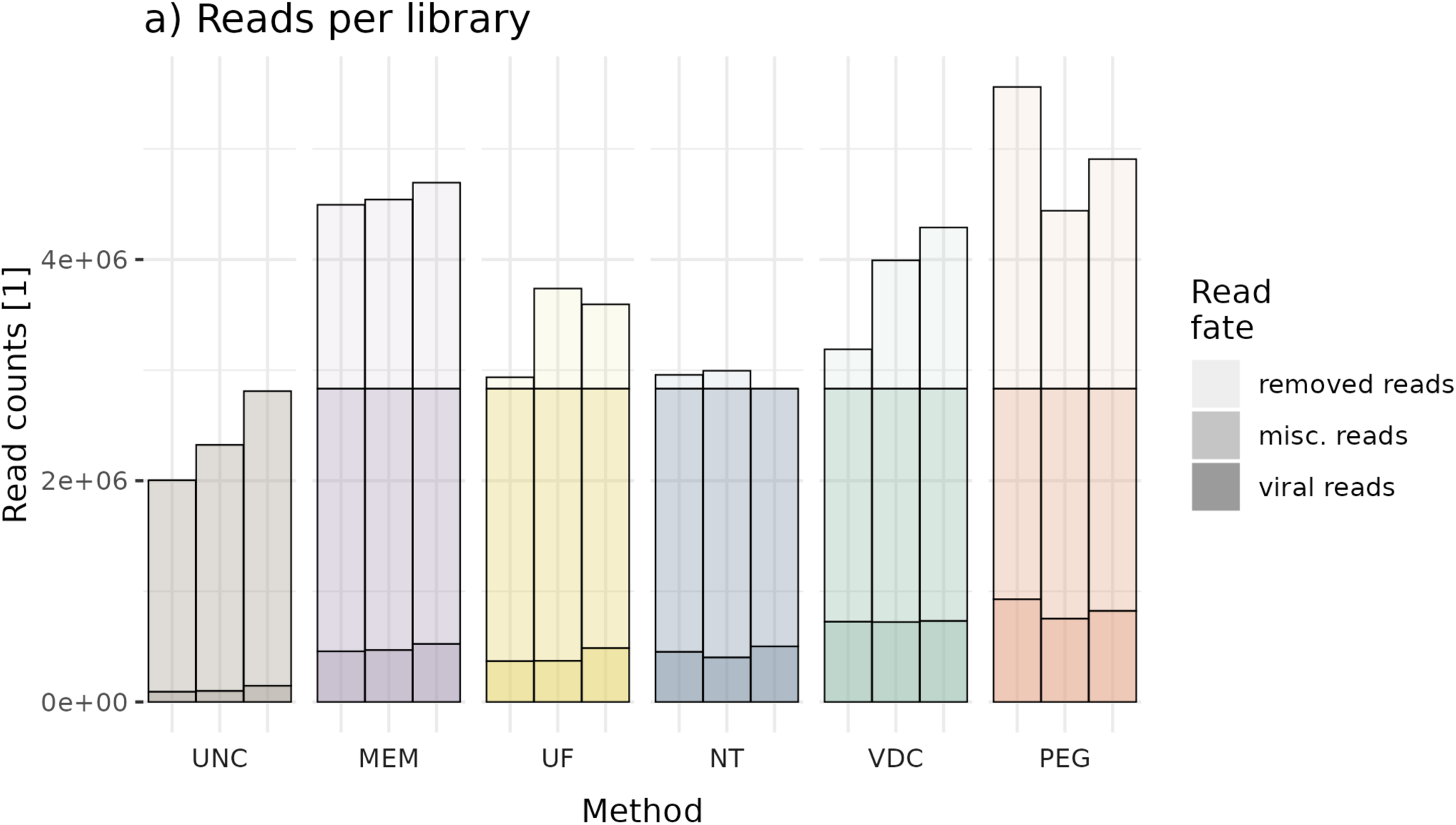
Read fates per library. Transparency levels indicate the read number at each stage. Most transparent bars (removed reads) indicate the raw library size, the second level (misc. reads) indicates the reads left over following down sampling, and the final level (viral reads) indicates reads that mapped to a virus.

**Supplemental Figure S5:**
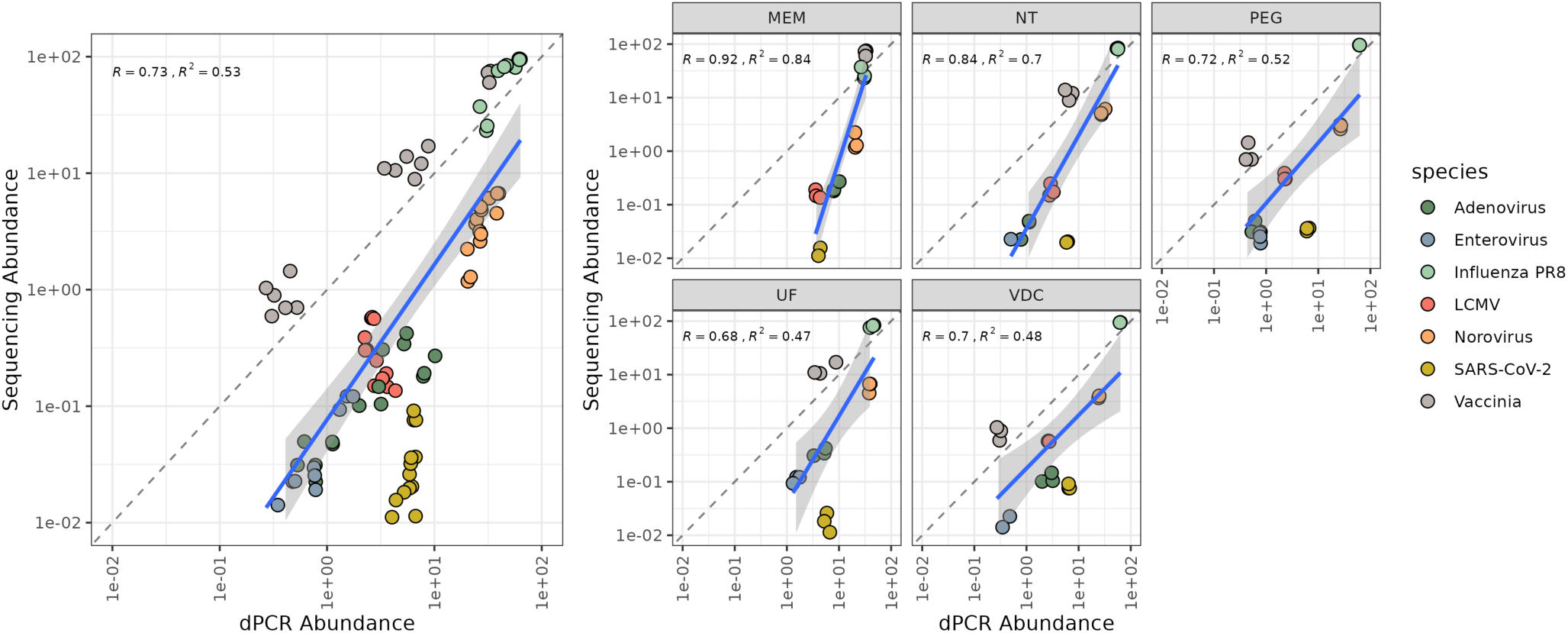
Comparison between dPCR and sequencing quantification. Virus abundance, read counts for sequencing data and concentration for dPCR data, normalized by sum of all target virus abundances that were measured by dPCR and present on the hybrid-capture panel, correlated with each other.

**Supplemental Figure S6.**
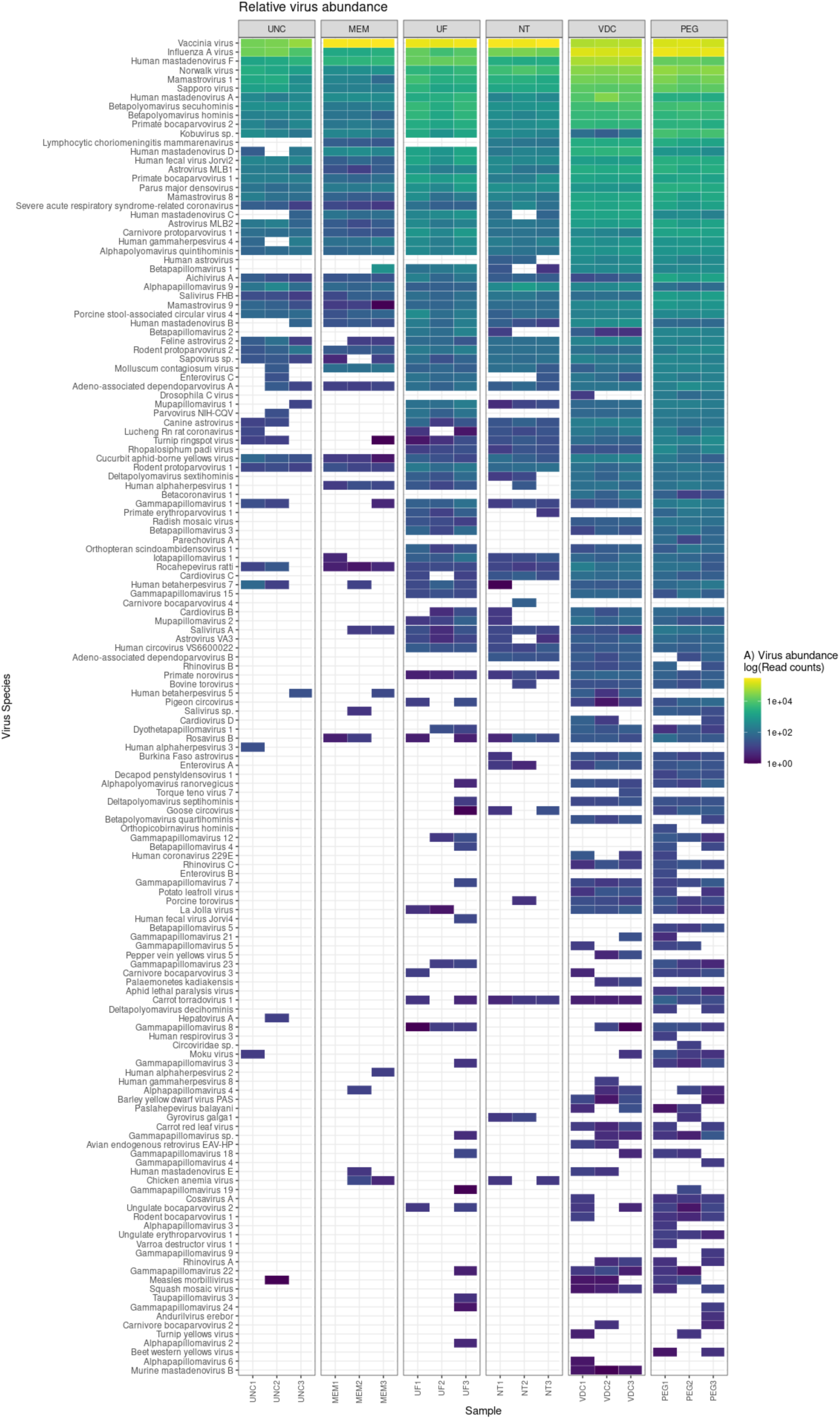
Viral Abundances with species names. Recreation of subpanel A of Figure 2 including the viral species names. All other aspects of the figure are the same.

**Supplemental Figure S7.**
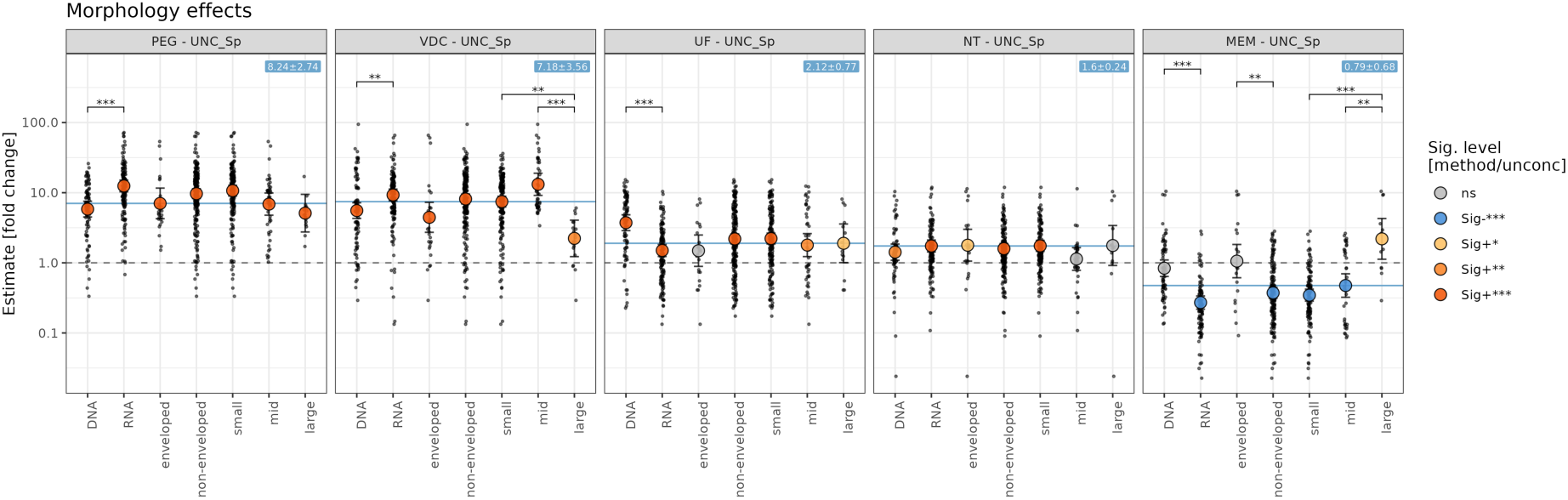
Modeling results of viral physical attributes. Model estimates for method enrichment performance across different groups of physical virus attributes collected at the genus level. Size classes were defined as small <75 nm, medium between 75 and 150 nm, and large >150 nm. Point color indicates significance of enrichment from the unconcentrated values after multiple testing correction, and error bars represent the 95% confidence interval. Significance brackets compare attribute enrichment effects within an attribute class (eg. size, nucleotide type, envelope status). Jitter points represent the raw data values.

**Supplemental Figure S8.**
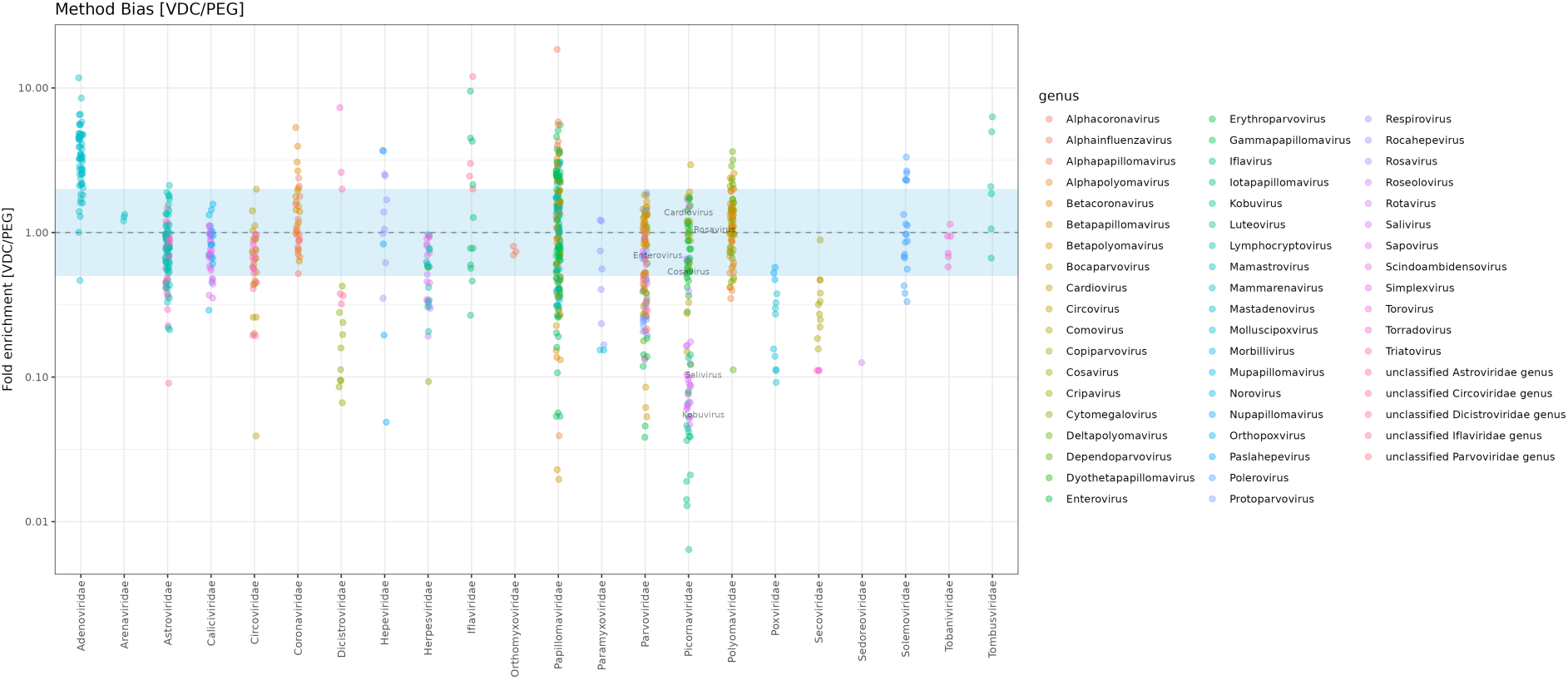
Read ratios of VDC to PEG. Each point is a read ratio of a viral strain detected in both PEG and VDC. Colors represent the genus the strain belongs to. Labelled are the genera of Pirconaviridae to highlight the genus-specific effect. Labels are vertically centered at the mean bias value for the genus.

**Supplemental Figure S9:**
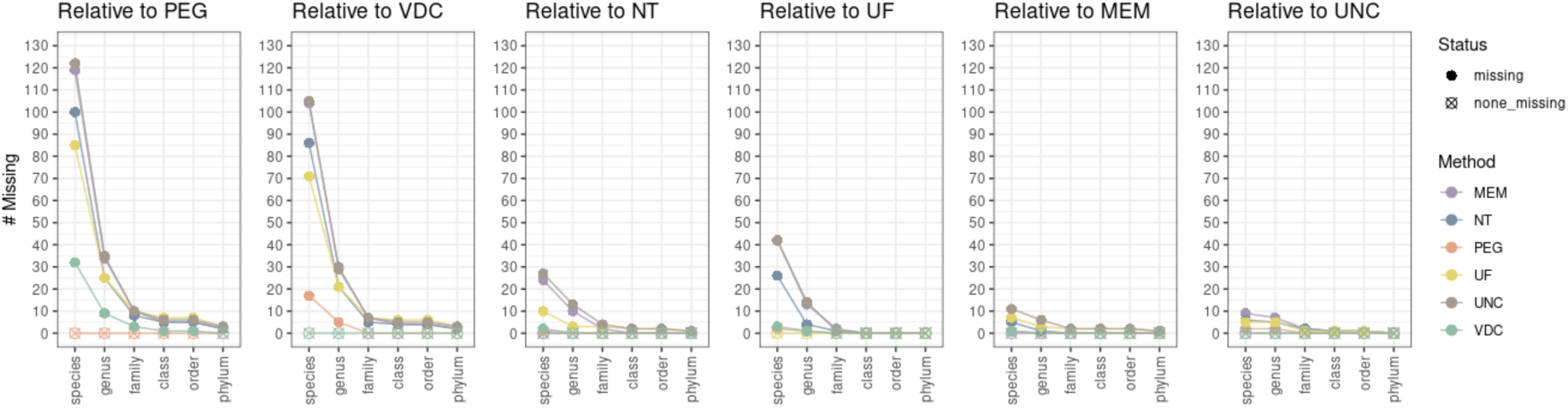
Missing taxa. Points show the number of taxa present in the reference method but missing in the comparison methods. For example, points in the first panel represent the number of taxa missed in VDC, NT, UF, MEM, or UNC but present in PEG samples. Color indicates the comparison method and reference method is indicated in the plot title.

## Supplemental Tables

**Supplemental Table S1:** Targets with dedicated dPCR assays, viral characteristics and information on method performance based on dPCR analysis in this study. Method performance was evaluated based on mean recovery for all targets except rotavirus or raw dPCR quantifications corrected for sample input for rotavirus. Best, best performance ± 1% mean recovery/normalized dPCR quantifications; good, ≥ 35% of best performing method; bad, < 35% of best performing method.

| Virus name | Type | Enveloped | Size (nm) | Nucleic Acid | Shape | Method performance |  |  |
| --- | --- | --- | --- | --- | --- | --- | --- | --- |
|  |  |  |  |  |  | Best | Good | Bad |
| Vaccinia Virus | Spike-in | Yes | 260 | ds DNA | Ellipse | NT, MEM | - | UF, VDC, PEG |
| Influenza-PR8 | Spike-in | Yes | 100 | ss(-) RNA | Ellipse | VDC | NT, PEG | UF, MEM |
| LCMV (Arm, Cl13; 1:1) | Spike-in | Yes | 100 | ss(-) RNA | Spherical | VDC | NT, PEG | MEM, UF |
| PhiX174 | Spike-in | No | 25 | ss(+) DNA | Icosahedral | VDC, PEG | UF | NT, MEM |
| Bacteriophage MS2 | Spike-in | No | 27 | ss(+) RNA | Icosahedral | VDC | PEG, NT, UF | MEM |
| SARS-CoV-2 | Endogenous | Yes | 100 | ss(+) RNA | Spherical | VDC | NT, PEG | UF, MEM |
| PMMoV | Endogenous | No | 18x312 | ss(+) RNA | Rod | VDC | NT, PEG | MEM, UF |
| Adenovirus (40, 41) | Endogenous | No | 80 | ds DNA | Polyhedral | VDC | MEM | NT, UF, PEG |
| Rotavirus A | Endogenous | No | 75 | ds RNA | Icosahedral | VDC | MEM, PEG, NT | UF |
| Enterovirus | Endogenous | No | 30 | ss(+) RNA | Icosahedral | NT, PEG, VDC | UF, MEM |  |
| Norovirus | Endogenous | No | 39 | ss(+) RNA | Icosahedral | VDC | NT, PEG | UF, MEM |

**Supplemental table S2.**
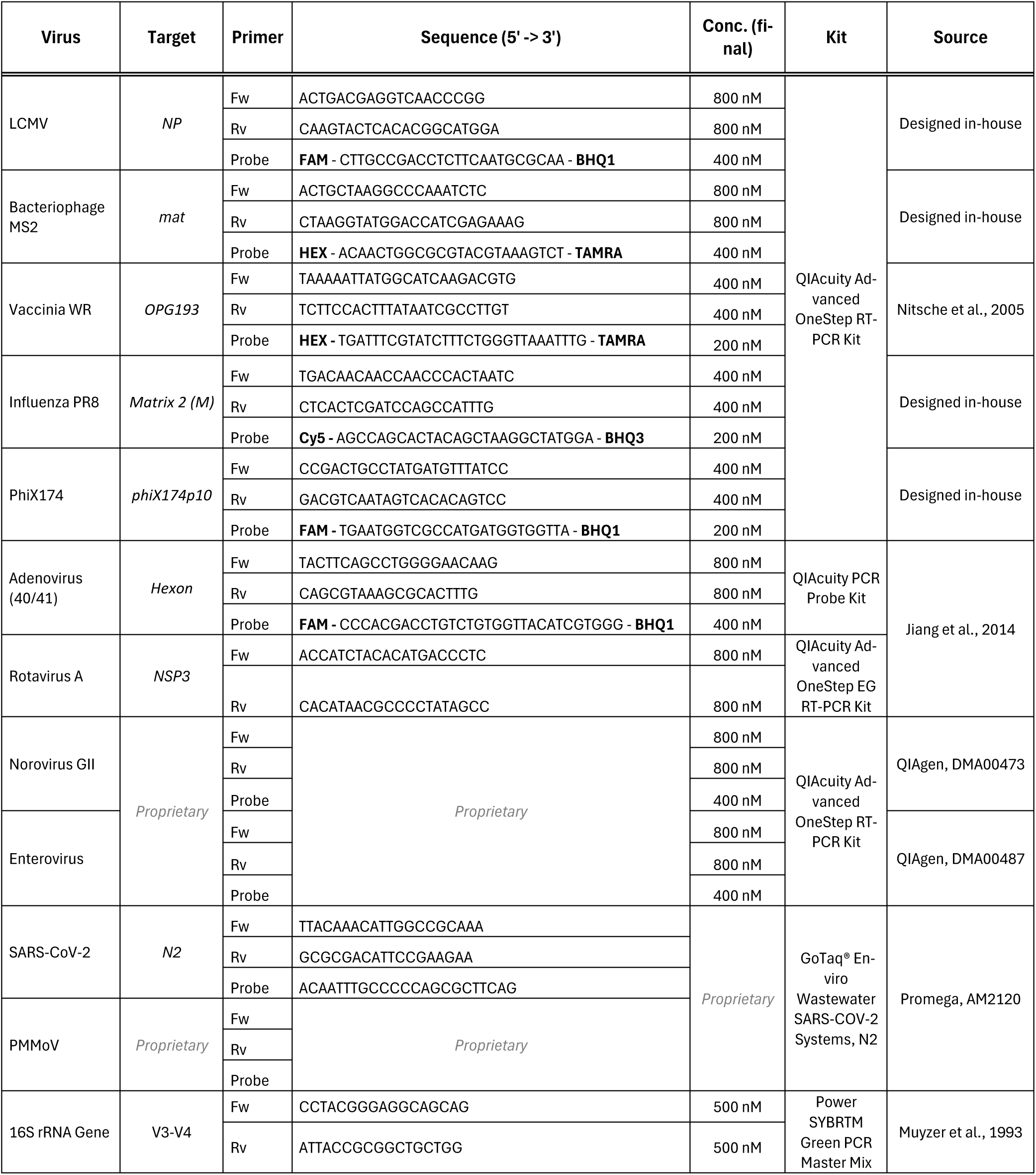
dPCR conditions and concentrations. Listed is the information for all PCR assays used in this study. Cycling conditions followed the standard recommendations for the respective kit used.

